# Electron cryomicroscopy observation of acyl carrier protein translocation in type I fungal fatty acid synthase

**DOI:** 10.1101/636167

**Authors:** Jennifer W. Lou, Kali R. Iyer, S. M. Naimul Hasan, Leah E. Cowen, Mohammad T. Mazhab-Jafari

## Abstract

During fatty acid biosynthesis, acyl carrier proteins (ACPs) from type I fungal fatty acid synthase (FAS) shuttle substrates and intermediates within a reaction chamber that hosts multiple spatially-fixed catalytic centers. A major challenge in understanding the mechanism of ACP-mediated substrate shuttling is experimental observation of its transient interaction landscape within the reaction chamber. Here, we have shown that ACP spatial distribution is sensitive to the presence of substrates in a catalytically inhibited state, which enables high-resolution investigation of the ACP-dependent conformational transitions within the enoyl reductase (ER) reaction site. In two fungal FASs with distinct ACP localization, the shuttling domain is targeted to the ketoacyl-synthase (KS) domain and away from other catalytic centers, such as acetyl-transferase (AT) and ER domains by steric blockage of the KS active site followed by addition of substrates. These studies strongly suggest that acylation of phosphopantetheine arm of ACP may be an integral part of the substrate shuttling mechanism in type I fungal FAS.

---

Type I fungal FAS forms a barrel-shaped complex composed of two reaction chambers. In *Saccharomyces cerevisiae*, the *FAS1* and *FAS2* genes produce a 230kDa β- and a 220kDa α-chain, respectively, that assembles into a heterododecamer of α_6_β_6_.^1,2^ Six β-chains form the walls of the barrel while a central wheel, made by the six α-chains, bisects the barrel into two chemically identical reaction chambers. Each chamber is formed by the central α-wheel and three β-chains around a C3 axis of symmetry. The two chambers are related by C2 symmetry, making the complex D3 symmetric (Supplementary Figure 1A). Therefore, each chamber has three complete sets of catalytic domains including three acyl-carrier protein (ACP) domains. ACP in type I fungal FAS is an 18kDa, eight helical domain with a four helical subdomain that is similar to the canonical type I metazoan and type II bacterial FAS ACP (herein refer to as canonical lobe) and an additional four helical subdomain (herein refer to as structural lobe). In the atomic resolution crystal structures of *S. cerevisiae* FAS, ACP is seen at the KS-binding site with both lobes of the domain contributing to the binding interface.^1,2^ ACP interacts twice with the KS domain during each catalytic cycle, unlike other catalytic sites where the mobile domain only interacts once^3^ (Supplementary Fig. 1A). Therefore, it is speculated that ACP interaction with other reaction sites is more transient. The canonical lobe is post-translationally modified with a phosphopantetheine moiety catalyzed by phosphopantetheinyl transferase (PPT) domain.^4^ This reaction creates holo-ACP, which can covalently bind substrates and reaction intermediates allowing the fungal FAS to carry out the multi-step synthesis of palmitoyl-coenzyme A.^3^ Except for the PPT domain, all catalytic centers face the interior of the chamber. Substrates are shuttled between the static reaction centers by the mobile ACP domain flexibly tethered at its N and C termini. A challenge in biophysical study of type I fungal FAS is experimental observation of the interaction landscape of the mobile ACP within the reaction chambers. In near-atomic resolution electron cryomicroscopy (cryoEM) maps of type I fungal and atomic-resolution cryoEM maps of type I bacterial FAS, ACP density is heterogenous as it samples multiple locations within the reaction chamber.^5–7^ Therefore methods that can modulate localization of ACP within the reaction chambers of fungal FAS, may improve ACP visualization in experimental cryoEM or X-ray crystallography density maps. Here, we have experimentally probed for the ability to redistribute ACP, by stalling catalysis at the KS site in two type I fungal FAS.

We used cryoEM to reconstruct *ab initio* ACP densities inside the reaction chambers of endogenous fungal FASs from *S. cerevisiae* and the opportunistic pathogen *Candida albicans* in the Apo and KS-stalled state, at 12 Å resolution, allowing localization of densities corresponding to this mobile domain (Supplementary Figure 2A-D). The *ab initio* ACP densities were generated using an ACP-less initial cryoEM density map that was generated from the ∼3 Å resolution atomic model of *S. cerevisiae* FAS^2^ with ACP atoms deleted and low-pass filtered to 30 Å. For simplicity, we call these maps ACP-*ab initio* (AAI) maps. The AAI maps were scaled relative to each other for comparison of ACP densities between Apo and KS-stalled reconstructions for each fungal species and are shown at identical resolution range (*i.e.* 30-12 Å) and threshold, unless otherwise stated. The AAI maps were then refined to high resolution (Supplementary Figure 2E, 3 and 4, Supplementary Table 1) to probe i) ability to resolve ACP helices and its phosphopantetheine moiety and ii) for ACP-mediated conformational transitions as discussed below.

In the Apo state, ACP density is strongest in proximity of the KS domain in *S. cerevisiae* FAS (Figure 1A and Supplementary Figure 2D and E) and allows for a complete tracing of its backbone atoms in the high resolution cryoEM map (Supplementary Figure 5A). In the Apo state of *C. albicans* (62 and 69% identity for β- and α-chains, relative to their respective *S. cerevisiae* chains), the ACP density is strongest in proximity of the ER domain, similarly allowing for complete tracing of the backbone atoms of this mobile domain at an alternate location (Figure 1B and Supplementary Figure 2D and E).

**Figure 1.**
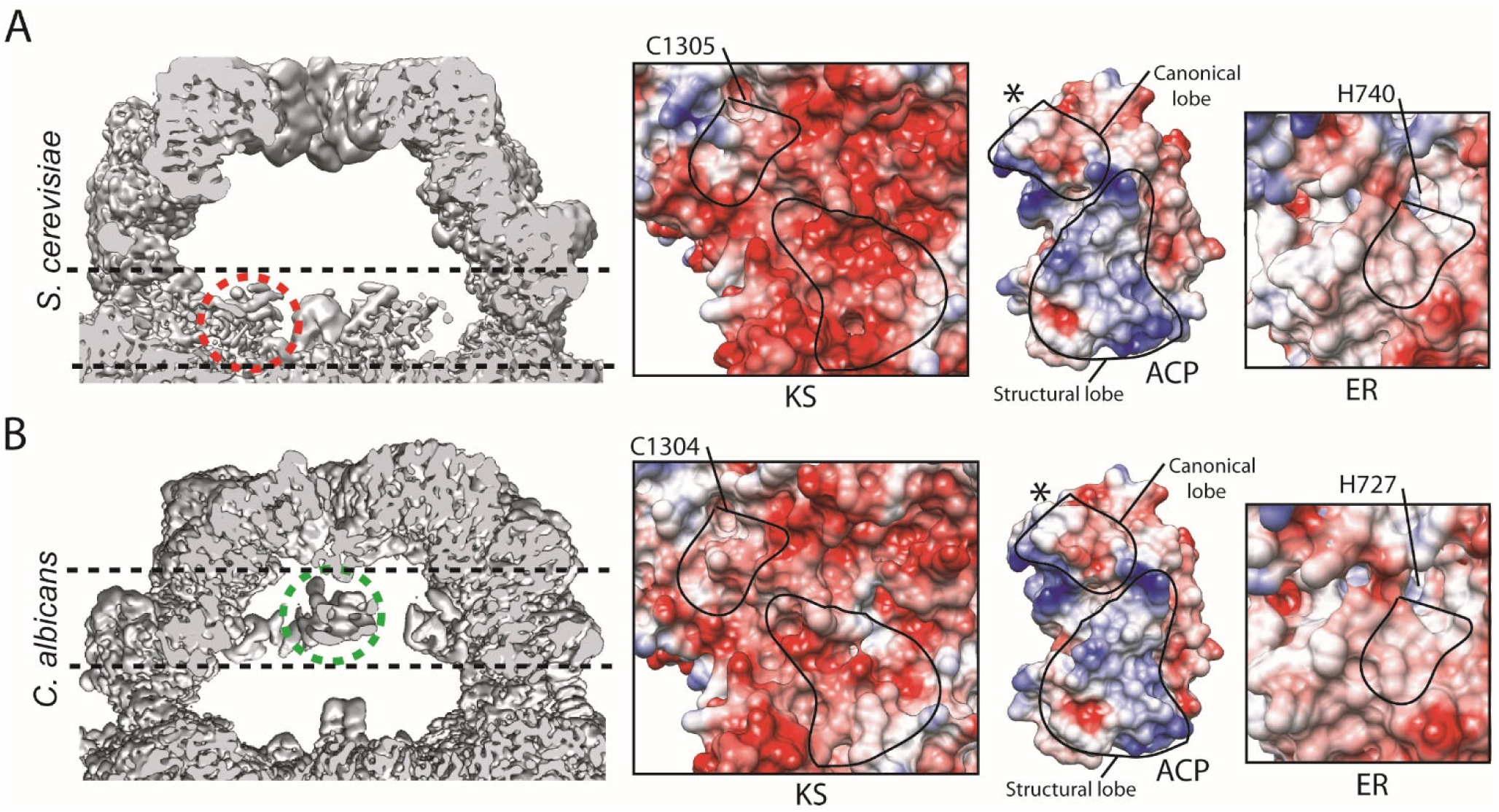
Different ACP localization within (A) *S. cerevisiae* and (B) *C. albicans* FAS in the Apo state. One ACP in each reaction chamber is highlighted with black dashed lines and red (*S. cerevisiae*) and green (*C. albicans*) circles. Coulombic surface coloring using Chimera^25^ for the interaction interfaces for KS-ACP- and ER-domains are shown. The interacting surface area for the portions of ACP’s structural and canonical lobes are encircled with continuous lines. The catalytic residues for KS and ER are highlighted and * represents the position of the phosphopantetheine arm of ACP.

Surface electrostatics have been previously speculated to be a steering force for ACP-mediated substrate shuttling based on molecular dynamic simulations.^8^ Examining the surface electrostatics of ACP-KS- and ER-domains shows a strong surface charge complementarity between the structural lobe of ACP with the KS domain in *S. cerevisiae* (Figure 1A). However, the negative surface charge on KS domain is weakened in *C. albicans* due to alteration of some of the acidic residues that form the interface with the structural lobe of ACP (Figure 1B, supplementary Figure 6A). A cryoEM map of a thermophilic fungal (*i.e. Chaetomium thermophilum*) type I FAS also observed ACP in proximity of the ER domain, albeit at a global resolution of 4.7 Å, which allowed for domain docking of ACP and assignment of partial density to the phosphopantetheine arm.^7^ Interestingly, this pattern of weakened negative surface charge is predicted to be preserved in *C. thermophilum* FAS based on sequence alignment (Supplementary Figure 6A). Weaker charge complementarity can partly explain why in this pathogenic fungal species, ACP is not primarily localized at the KS in the Apo state. There are no significant alterations in the surface electrostatics of the ACP binding site in the ER domain between the *S. cerevisiae* and *C. albicans* (Figure 1) and residues lining the ACP binding site on the ER domain are mostly conserved between the two species studied here and the *C. thermophilum* (Supplementary Figure 6B). Interestingly, unlike the KS-binding site where both canonical and structural lobes of ACP contribute to the surface area of the binding interface, the interface of ACP at the ER mainly involves the canonical lobe (Supplementary Figure 5B). This observation can explain the worse local resolution seen for the ACP of *C. albicans* localized at the ER compared to that in *S. cerevisiae* at the KS domain in the Apo states (Supplementary Figure 2E). It is possible that the transient interactions of ACP that are mainly driven by its phosphopantetheine-containing canonical lobe are more prone to chemical and structural alterations.

In both fungal FASs, weaker densities can also be seen proximal to other catalytic centers in the Apo state. In *S. cerevisiae* FAS, second strongest ACP density is observed at lower resolution and less stringent threshold of the AAI map that shows ACP near the apical region of the reaction chamber and facing the AT catalytic site (Figure 2A, Supplementary Figure 2D). In the Apo state of *C. albicans* FAS, the second strongest density is in proximity of the KS domain that is only observable at lower resolution and threshold of the AAI cryoEM density map (Figure 2B, Supplementary Figure 2D). These ACP densities do not refine to high-resolution to characterize and model their respective binding interfaces in the Apo states of the two fungal FASs (Supplementary Figure 2E), most likely due to a lower occupancy and less stable interaction compared to that of ACP with the KS and ER domains in *S. cerevisiae* and *C. albicans*, respectively.

**Figure 2.**
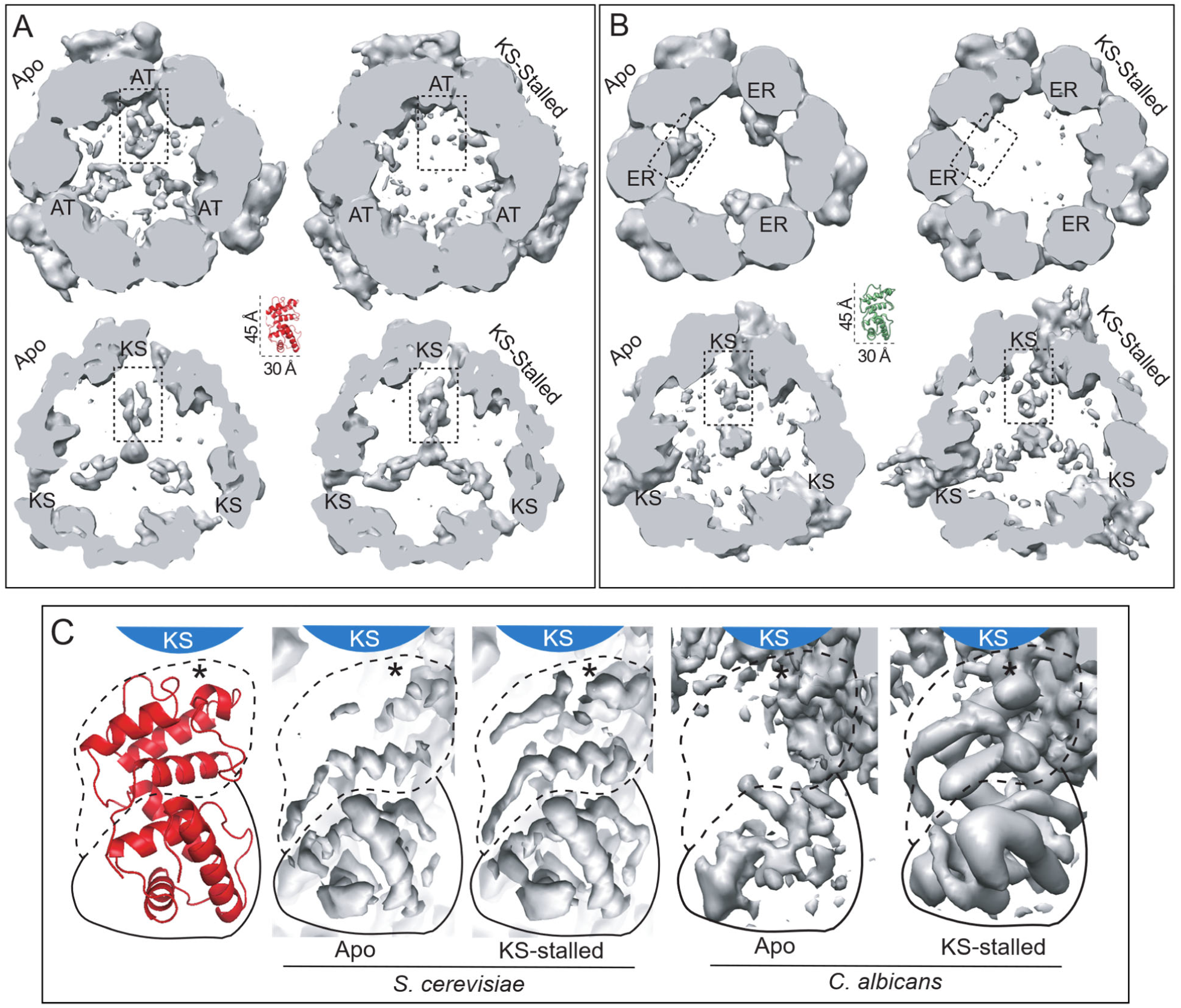
ACP distribution and translocation in *S. cerevisiae* and *C. albicans* FAS. Slices through the reaction chamber near (A) AT- and KS-domains for *S. cerevisiae*, and (B) ER- and KS-domains for *C. albicans* in the Apo and KS-stalled states. See supplementary Figure 2D for position of slices. Slices for each catalytic centre from *Ab initio* maps are shown at identical resolution range and threshold in panels (A) and (B). Model of ACP shown to scale at the center of panels A and B in red and green for *S. cerevisiae* and *C. albicans*, respectively. (C) High-resolution refined maps of FAS showing improved high-resolution features of the structural (solid line) and canonical (dashed line) lobes of ACP in proximity of the KS catalytic cavity in the KS-stalled state for each fungal species. Atomic model of ACP with the same orientation as the maps is shown to the left in red. * represent the position of the phosphopantetheine arm.

In type II bacterial FAS, where ACP is only composed of the canonical lobe, substrate loading state can modulate the affinity profile of a panel of ACP mutants toward KS protein and active-site specific cross-linkers have proven to be valuable tools to trap transiently formed ACP complexes with other catalytic proteins such as ER and dehydratase (DH).^9–12^ We therefore hypothesized that the loading-state of ACP may modulate its interaction with the catalytic centers in the context of fully assembled type I fungal FAS and allow for redistribution of ACP toward a specific reaction site. Both endogenously purified enzymes are catalytically active (*i.e.* contain holo-ACP) and are sensitive to cerulenin, a covalent KS-inhibitor, as judged by monitoring NADPH oxidation at 340nm (Supplementary Figure 1B). In the crystal structures of cerulenin-inhibited *S. cerevisiae* FAS, partial density could be seen for the inhibitor, which allowed model building for a portion of the compound.^13,14^ In both our activity assays and cryoEM samples, the FAS enzymes were pre-incubated with cerulenin prior to addition of substrates acetyl- and malonyl-CoA and NADPH to ensure KS site was inhibited prior to the start of the reaction (*i.e.* KS-stalled state). In our high-resolution cryoEM maps, we can see appearance of additional densities immediately above the catalytic cysteine of the KS-domain in the KS-stalled state suggesting covalent linkage between cerulenin and the KS catalytic residue (Supplementary Figure 4C), which confirms our activity assay demonstrating inhibition of catalysis at the ketoacyl synthesis step (Supplementary Figure 1).

Stalling catalysis at the KS domain drastically alters the ACP densities inside the reaction chamber of both fungal FASs, as judged by improved ACP density near KS domain and weakened ACP density by the AT- and ER-domains in *S. cerevisiae* and *C. albicans* FASs, respectively (Figure 2A and B). In both FASs in the Apo state, the structural lobe of ACP has stronger and better resolved densities compared to the canonical lobe in the KS-binding site, as observed in the high-resolution refined maps (Figure 2C), suggesting a more stable interaction between the former and the KS-domain in the absence of substrates. In both species, the major improvement in cryoEM density of ACP at the stalled KS domain is observed at its canonical lobe (Figure 2C). These observations suggest a bimodal interaction between ACP and the KS domain, one mediated by the ACP structural lobe and the other by its canonical lobe, that may be modulated independently by the state of the ACP- and KS-catalytic sites. In the KS-stalled state, ACP is presumably acylated with acetyl or malonyl as it can not transfer the starting (*i.e.* acetyl loaded by AT domain) and elongating (*i.e.* malonyl loaded by MPT domain) substrates to KS from its phosphopantetheine arm. However, in our cryoEM maps of *S. cerevisiae* (where ACP density is better resolved in proximity of the KS domain relative to *C. albicans*), density for the phosphopantetheine arm at the KS-catalytic cavity can only be seen until the phosphate group (Supplementary Figure S7), likely due to conformational heterogeneity of the moiety combined with ACP’s transient interaction with KS. Therefore, we can not experimentally observe a complete density of the acylated phosphopantetheine and can only infer its acylated state based on the catalytic competency of the enzyme and its inhibition with cerulenin (Supplementary Figure 1B and 4C). The observation that stalling catalysis at one site (*i.e.* KS) can affect ACP localization at other sites (*i.e.* AT and ER) in the presence of substrates within the context of the fully assembled reaction chamber strongly suggests that substrate loading state of the mobile domain may be a contributing factor in ACP-mediated substrate shuttling in type I fungal FAS.

With the ability to observe ACP at the ER and redirect this mobile domain to the KS, we focused on conformational differences within the ER catalytic region of *C. albicans* FAS. This site within *C. albicans* enzyme represented the most pronounced alteration in ACP localization upon stalling catalysis at the KS-domain (Supplementary Figure 2D and E). The ER domain shows the highest sequence and structural divergence between type I human and fungal FAS and is an ideal drug target for infectious diseases including *Candida* infections in immune compromised patients. Unlike human FAS,^15^ the fungal ER domain uses a flavin mononucleotide (FMN) co-factor for the conversion of enoyl-ACP to saturated acyl-ACP. The FMN is then replenished by NADPH that can bind a pocket accessible from outside the reaction chamber.^16^ The histidine residue adjacent to FMN is proposed to catalyze proton transfer based on homology to FMN-dependent bacterial protein FabK.^17,18^ The loop hosting the catalytic histidine is highly conserved between *S. cerevisiae* and *C. albicans* (Figure 3A).

**Figure 3.**
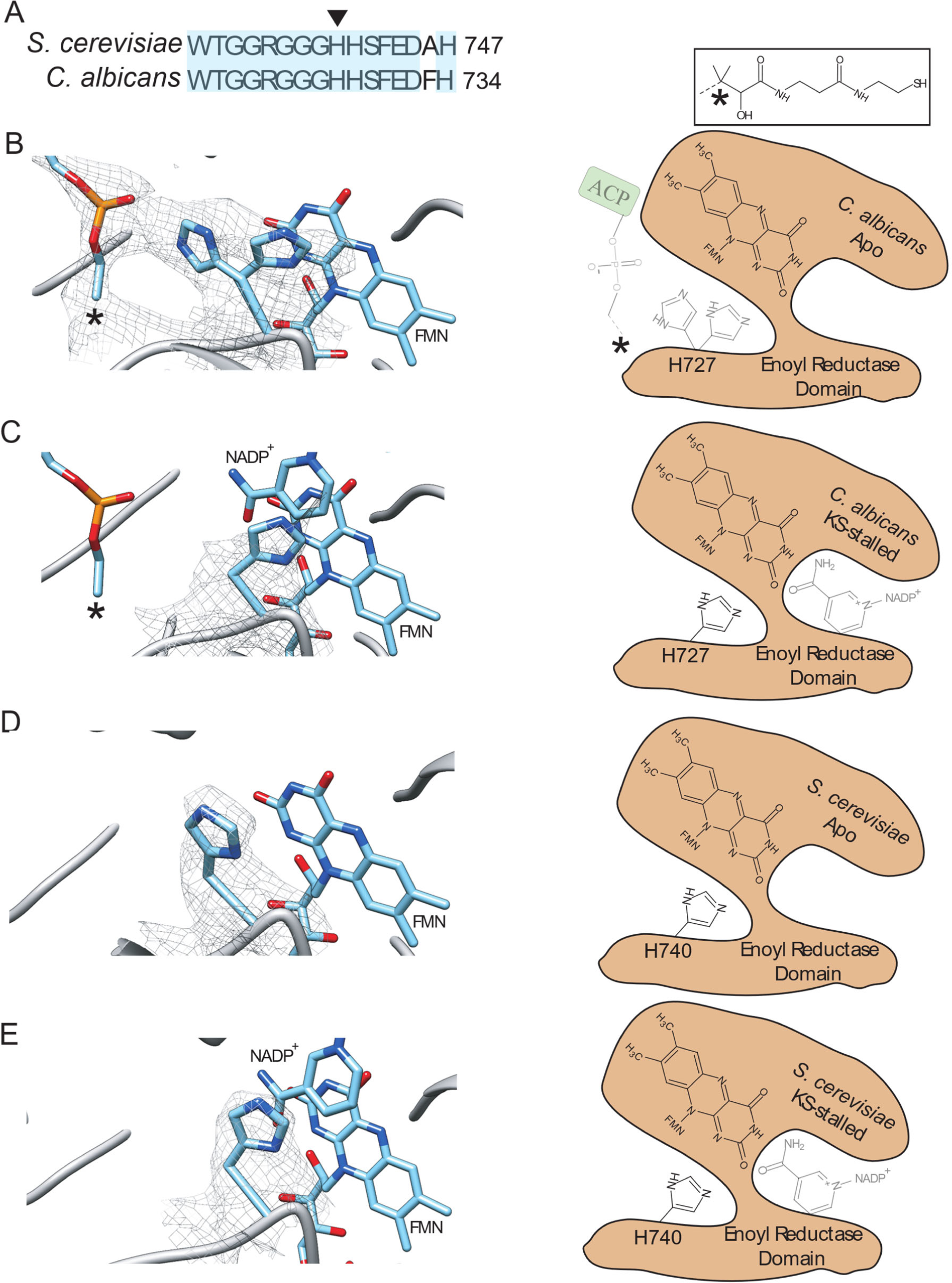
Conformational rearrangement of catalytic histidine of the ER-domain upon ACP-binding. (A) Sequence alignment of the ER catalytic loop between the two fugal FAS with the conserved region highlighted in cyan. Catalytic histidine shown with arrow head. cryoEM densities are shown within 3.5 Å of the of catalytic histidine and the atoms of the phosphopantetheine arm as shown in the (B) Apo- and (C) KS-stalled states of *C. albicans*. (D) and (E) are cryoEM densities of the catalytic histidine, shown within 3.5 Å in the Apo- and KS-stalled state of *S. cerevisiae* FAS, respectively. For clarity, densities of FMN and NADPH are not shown. The schematic diagrams of the ER catalytic site are shown to the right for each panel. * represents the unmodeled portion of the phosphopantetheine prosthetic group. Transparent schematics represent partial occupancy.

Examining the ER active site in *C. albicans* Apo state where ACP is partially bound, we can clearly see the density of the phosphopantetheine arm until the quaternary carbon atom of the moiety (Figure 3B). Interestingly, the catalytic histidine can be modeled with two side chain orientations, one facing FMN and the other facing the phosphate group of the phosphopantetheine arm. When ACP is shifted away from the ER domain in the KS-stalled state, the catalytic histidine is mainly in one conformation facing FMN (Figure 3C). Weak density can be seen for the NADPH in proximity of FMN in the KS-stalled state, indicating partial/transient binding that does not clash with either of the histidine side chain conformations seen here (Supplementary Figure 8A). Interestingly, in the *S. cerevisiae* Apo and KS-stalled states, where no ACP density can be seen by the ER domain, the catalytic histidine side chain is in a single conformation facing FMN (Figure 3D and E). Weak NADPH density can also be seen in proximity of FMN in the KS-stalled *S. cerevisiae* FAS (Supplementary Figure 8B). During catalysis, the enoyl group of the elongating fatty acid chain should come in proximity of the catalytic histidine to enable the reduction reaction. Based on these structures, the catalytic histidine in the ER domain of type I fungal FAS may sample alternative conformations to form a proton transfer bridge between FMN and the enoyl functional group of the elongating fatty acid chain when ACP is bound.

Structural knowledge on the interaction landscape of ACP within type I FAS and understanding the mechanism of ACP-mediated substrate shuttling are highly sought after in antibiotic development^19–21^ and biofuel production efforts.^22–24^ This study demonstrates that ACP interaction with a catalytic center can be modulated by stalling catalysis at another reaction site, suggesting that the loading state of ACP’s phosphopantetheine arm may have at least a partial role in determination of ACP localization within the reaction chambers of type I fungal FAS. Future structural studies should systematically investigate ACP distribution upon inhibition of each reaction site in FAS (Supplementary Figure 1A) using site-specific inhibitors or point mutations in recombinantly expressed enzymes.

## Acknowledgements

Images were collected at the Toronto High Resolution High Throughput cryo-EM facility, supported by the Canada Foundation for Innovation and Ontario Research Fund. We thank Dr. John L. Rubinstein for kindly providing the *S. cerevisiae* yeast strains and plasmids, access to the FEI Tecnai F20 electron microscope, and critical review of this manuscript. We thank Dr. Zhongle Liu for kindly providing *C. albicans* plasmids, and Dr. Jaideep Mallick for assistance with *C. albicans* strain validation.

This work is supported by Princess Margaret Cancer Research Institute and Princess Margaret Cancer Foundation (PMCF). JWL was supported by Natural Sciences and Engineering Research Council of Canada (NSERC). MTMJ and SMNH were supported by PMCF. LEC is supported by a Canada Research Chair in Microbial Genomics and Infectious Disease and the Canadian Institutes of Health Research Foundation Grant (FDN-154288).

## Author contributions

JWL and MTM-J conceived and designed the research. KRI prepared genetically modified *C. albicans* cells. LWC, supervised preparation of modified *C. albicans* strain. JWL prepared the genetically modified *S. cerevisiae* cells and performed protein purification and functional assays. JWL and SMNH prepared EM and cryoEM samples. MTM-J collected EM and cryoEM images using F20 Tecnai electron microscope. JWL and MTM-J analyzed the images and generated cryoEM maps. JWL and MTM-J created models. JWL and MTM-J interpreted the data. JWL, LWC, and MTM-J prepared the figures and wrote the paper.

