## Supplementary methods, Tables, and figures for "Electron cryomicroscopy observation of acyl carrier protein translocation in type I fungal fatty acid synthase"

### Supplementary Information:

**Supplementary Table 3**…………………………………....……………7

**Supplementary Figure 1**………………………………………….…..8

**Supplementary Figure 2**…………………………..…...………..10

**Supplementary Figure 3**……………………………….……………..12

**Supplementary Figure 4**……………………………….……………..13

**Supplementary Figure 5**…………………….……….....………..15

**Supplementary Figure 6**…………………….………………………..16

**Supplementary Figure 7**…………………….………………………..18

**Supplementary Figure 8**…………………….………………………..19

### Materials and Methods:

#### Yeast strains and protein purification:

The genomic DNA of haploid protease deficient *S. cerevisiae* strain BJ2168 (MATa leu2 trp1 ura3-52 prb1-1122 pep4-3 prc1-407 gal2) was modified by homologous recombination method[1,2] to encode for a 3×FLAG tag at the C terminus of *FAS1* gene. To construct the modified yeast strain (JWL01) expressing FLAG-tagged FAS1, the 3xFLAG-URA3 cassette was amplified from pJT1 plasmid[3] using primers containing 50 bp homology to the FAS1 C terminus (SI Table S2). The amplified fragment was used to transform the yeast cells by a lithium acetate-based method[1]. Transformants were selected on synthetic media (SD) uracil-dropout plates and confirmed with PCR using a forward primer chosen from within the *FAS1* gene and a reverse primer chosen from within the inserted cassette (SI Table S2).

To construct *C. albicans* yeast strain CaLC5425, the 6xHIS-3xFLAG-HIS cassette was amplified from pLC1085 plasmid[4] using the primers oLC6912 and oLC6913 which contained 70 bp homology to *C. albicans* FAS1 C-terminus (SI Table S3). The amplicon was then transformed into SN95 (ura3::imm434::URA3/ura3iro1IRO1/iro1his1his1arg4/arg4) strain. Cells were plated on SD+ARG plates and HIS+ transformants were PCR tested for correct integration of 6xHIS-3xFLAG at the C terminus of FAS1 using the primer pairs oLC6914/oLC6915 and oLC6916/oLC6917 (SI Table S3).

To purify endogenous yeast FAS complexes, 1L *S. cerevisiae* yeast in YPD media were grown in 4L flasks in an Innova 42 shaker (New Brunswick) at 25°C until $OD_{660nm}$ ~ 2.0, or mid-exponential phase. *C. albicans* cells were grown in 600mL YPD medium in 2L flasks shaking (Thermo Electron) at 30°C until an $OD_{600nm}$ ~ 3.0 was reached. Cells were harvested by centrifugation at 4,000 rpm for 10 min, flash frozen in liquid nitrogen and stored at -80C. For purification, cells were resuspended in lysis buffer (200 mM potassium phosphate pH 7.4, 10 mM EDTA, 0.5 mM PMSF, 50 mM β-glycerophosphate, 10 mM NaF, 5 mM aminocaproic acid, 1 mM benzamidine). Lysis was achieved by mechanical disruption at ice-water temperature using a BeadBeater (BIoSpec Products) with bead beating for 30s and resting for 60s for a total of 270s using 0.5 mm diameter glass beads. The lysate was cleared by an initial centrifugation at 4,000 g for 10 min followed by ultracentrifugation at 110,000 g for 30 min. The cleared lysate was filtered using a 0.22 um pore size filter and loaded onto a pre-equilibrated column containing 500 µL of anti-FLAG M2 affinity resin (Sigma-Aldrich). The column was washed 10 times with one column volume of lysis buffer and 10 times with one column volume elution buffer (50 mM Tris-HCl pH 7.4, 150 mM NaCl) and eluted with 3 column volumes of

150 ug/mL 3×FLAG peptide (GenScript) in elution buffer. 1mM DTT was added to the purified sample after elution. Purity was assessed by Coomassie Blue stained 8-15% gradient SDS-PAGE.

**Activity assays:**

Activity of purified FAS complexes at 0.2 mg/ml were measured through a 1 cm path length corrected spectrophotometric assay monitoring the level of NADPH (Sigma-Aldrich) at 340 nm. The assay mixture contained 100 mM potassium phosphate pH 7.4, 1 mM EDTA, 0.2 mM acetyl-CoA (Sigma-Aldrich), 0.7 mM NADPH, 1 mM DTT in a 100 µl reaction volume. The absorbance at 340 nm was measured for 3 minutes, after which the reaction was initiated by the addition of 30 nmol malonyl-CoA (Sigma-Aldrich) and monitored for 15 minutes at 25°C. To test the effect of the inhibitor on the enzymatic activity, FAS was incubated with cerulenin (Sigma-Aldrich) at room temperature for 1 h prior to the activity assay. For the positive control reactions (absence of cerulenin), DMSO was added to 1% v/v to match the highest DMSO concentration tested in the inhibitory reactions (*i.e.* 100 µM cerulenin). Each curve is normalized against its first point of measurement after addition of malonyl-CoA.

**Electron cryomicroscopy:**

To prepare samples for imaging by electron microscopy, freshly purified FAS complexes were concentrated to 4 mg/mL using Amicon Ultra centrifugal filters (Millipore Sigma) before applying 3 µl onto the nanofabricated holey sputtered gold grid[5] with a hole size of ~2 µm. Grid freezing was done in Vitrobot Mark IV (FEI) with 3 sec blotting at 4°C, 100% humidity and using liquid ethane at liquid nitrogen temperature. For KS-inhibited state, purified FAS at 0.5 mg/ml was incubated with 25 µM of cerulenin for 1h at room temperature. The sample was then concentrated to 10 mg/ml followed by addition of acetyl-CoA, malonyl-CoA, and NADPH pH-adjusted mixture. The final concentrations were 4 mg/ml FAS, 1mM acetyl-CoA, 3 mM NADPH, and 1.4 mM malonyl-CoA. Freezing was done as with the Apo sample.

Cryo-grids were screened for ice-thickness, particle distribution, and sample behaviour with a FEI Tecnai F20 field emission electron microscope equipped with a Gatan K2 summit direct detector device (DDD) camera. Images were acquired in counting mode with 1.45 Å/pixel, 2 frames/s for 15 s, and an exposure rate of 1.2 $e^{-}/Å^{2}$/frame. Data collection was done with Titan Krios G3 electron microscope. See SI table S1 for details on data collection from Titan Krios electron microscope.

**Image processing and model building:**

Image processing and statistics can be found in SI table S1. All motion corrections were done with Alignframe_lmbfgs and Alignpart_lmbfgs[6] that are implemented in cryoSPARC V2[7]. Particle picking, 2D classification and refinement were also done with cryoSPARC V2. For generating ACP densities inside the reaction chamber, a full FAS complex was modeled using PDB: 2UV8[8] followed by deletion of all ACP atoms. The modified model was converted to a cryoEM density map using Chimera[9] and low-pass filtered to 30Å. *Ab initio* ACP densities were generated using this modified ACP-less map as the initial model with D3 symmetry imposed. All *ab initio* maps were set to a maximum resolution of 12Å (resolution range 30 to 12Å, in cryoSPARC V1). These maps are shown and referred to in figures 2A&B of the main text. All subsequent high-resolution refinements were done with the corresponding maps from *ab initio* reconstructions with ACP-less initial FAS models and D3 symmetry applied. These high-resolution refinements were done with cryoSPARC V2 nonuniform (NU-) refinement algorithm and the filtered maps were used for model building. In all NU-refinement, the dynamic mask threshold was set to 0.05 (range 0-1, default = 0.2) for better masking and refinement of ACP densities. The Fourier shell correlation (FSC) curves were calculated using the independently refined half maps with resolution assessed at 0.143 after correcting for the effects of masking maps. Unless otherwise stated high-resolution filtered maps (from NU-refinement) are shown in the figures. Scaling and thresholding of different maps were done in Chimera[9] using 'vop' command and volume viewer tool. Model building was done using the high-resolution locally-filtered nonuniform maps and PDB 2UV8[8] as initial model for *S. cerevisiae* FAS. For *C. albicans* FAS, a homology model was built with SWISS-MODEL[10] using PDB 2UV8[8] as template and subsequently refined. In both FAS systems, the quality of the ACP density did not allow for confident building of complete atomic models for this mobile domain. Therefore, polyalanine models of ACP were generated. Refinement was done with Phenix[11,12] real space refinement tool and manual model building was done with Coot[13]. Validation statistics were generated using Molprobity[14] and EMringer[15] software implemented within Phenix. All visualizations are done with Chimera[9] and PyMol[16]. Surface electrostatic potentials for the KS and ER-domains were estimated using the refined atomic models and Coulombic surface coloring tool in Chimera. For surface electrostatics of ACP, homology models were made for both fungal species with SWISS MODEL[10] using PDB: 2UV8 as template and used for Coulombic surface coloring.

**Supplementary Table 1)** Data collection and statistics.

| Data Collection | S. cerevisiae FAS (Apo) | S. cerevisiae FAS (KS-stalled) | C. albicans FAS (Apo) | C. albicans FAS (KS-stalled) |
|---|---|---|---|---|
| Microscope | Titan Krios G3 | Titan Krios G3 | Titan Krios G3 | Titan Krios G3 |
| Camera | FEI Falcon 3EC | FEI Falcon 3EC | FEI Falcon 3EC | FEI Falcon 3EC |
| Voltage | 300 kV | 300 kV | 300 kV | 300 kV |
| Magnification | 75,000× | 75,000× | 75,000× | 75,000× |
| Pixel size | 1.06 Å | 1.06 Å | 1.06 Å | 1.06 Å |
| Exposure | 43 electrons/Å$^2$ | 43 electrons/Å$^2$ | 43 electrons/Å$^2$ | 43 electrons/Å$^2$ |
| Exposure rate | 0.8 Electron/Å$^2$/Sec | 0.8 Electron/Å$^2$/Sec | 0.8 Electron/Å$^2$/Sec | 0.8 Electron/Å$^2$/Sec |
| Number of Frames | 30 | 30 | 30 | 30 |
| Defocus Range | 0.6-2.5μm | 0.6-2.5μm | 0.6-2.5μm | 0.6-2.5μm |
| **Image Processing** | | | | |
| Frame motion correction | Alignframe_lmbfgs | Alignframe_lmbfgs | Alignframe_lmbfgs | Alignframe_lmbfgs |
| CTF estimation | CTFFIND4 | CTFFIND4 | CTFFIND4 | CTFFIND4 |
| CTF cutoff | 5 Å | 5 Å | 5 Å | 5 Å |
| Particle picking software | cryoSPARC | cryoSPARC | cryoSPARC | cryoSPARC |
| Micrographs Used | 4,075 | 4,056 | 1,123 | 1,310 |
| Particle image motion correction | Alignpart_lmbfgs | Alignpart_lmbfgs | Alignpart_lmbfgs | Alignpart_lmbfgs |
| Particles Contributed | 637,823 | 594,818 | 92,958 | 24,417 |
| Reconstruction Software | cryoSPARC | cryoSPARC | cryoSPARC | cryoSPARC |
| Symmetry Applied | D3 | D3 | D3 | D3 |
| Global Resolution (FSC = 0.143) | 2.9 | 2.8 | 2.8 | 3.3 |
| **Model Building** | | | | |
| Modeling Software | Coot, Phenix | Coot, Phenix | Coot, Phenix | Coot, Phenix |
| Number of Residues build | 3,647 | 3,647 | 3,376 | 3,376 |
| RMS (Bond) | 0.006 | 0.008 | 0.013 | 0.006 |
| RMS (Angles) | 1.13 | 1.2 | 1.44 | 1.17 |
| Ramachandaran Outliers | 0.03% | 0.03% | 0.06% | 0.09% |
| C-beta outliers | 0.00% | 0.00% | 0.00% | 0.00% |
| Rotamer Outliers | 0.22% | 0.48% | 0.93% | 0.20% |
| Clashscore | 5.79 | 5.99 | 6.2 | 4.5 |
| MolProbity Score | 1.57 | 1.62 | 1.8 | 1.56 |
| EMRinger Score | 1.95 | 2.06 | 3.17 | 3.11 |
| PDB ID | XXXX | XXXX | XXXX | XXXX |

**Supplementary Table 2)** Primers for tagging 3' end of *FAS1* gene in *S. cerevisiae*.

| Name | Sequence (5' -> 3') |
|---|---|
| Homologous recombination F | CCGAACCTATCAAGGAAATCATCGACAACTGGGAAAAGTATGAACAATCCGACTACAAAGACCATGACGG |
| Homologous recombination R | CAGGAGTTTCAAAGTTAAATATTTCTTACGGTTATATAATCACTTAAGAAAATATCATCGATGAATTCGAGCTCG |
| Confirmation F | GAAGGTTGCTAGATTGGCCG |
| Confirmation R | GAGCGACCTCATACTATACC |

**Supplementary Table 3)** Primers for tagging 3' end of *FAS1* gene in *C. albicans*.

| Name | Sequence (5' -> 3') |
|---|---|
| oLC6912 | CCAATCAGTTTATGATTTGACTAAATCGGAAAAAATCAAGAGTATTTTAGATAACTGGGAACAATACGAAGGTCGACGGATCCCC |
| oLC6913 | ATAACAGGTCCTTTAAATAGCAAGTAAATAGAAATTTTATACATTTATTTATTATCTATACATTCTAAGTTCGATGAATTCGAGCTCG |
| oLC6914 | TGCTGTCATTCCATTGAAGG |
| oLC6915 | TAACTTCTGTCTCCTCATCCTC |
| oLC6916 | TTTAAAGTCAATAGGCATTCTCG |
| oLC6917 | AGCCCAACTGGTAAAAAGCA |

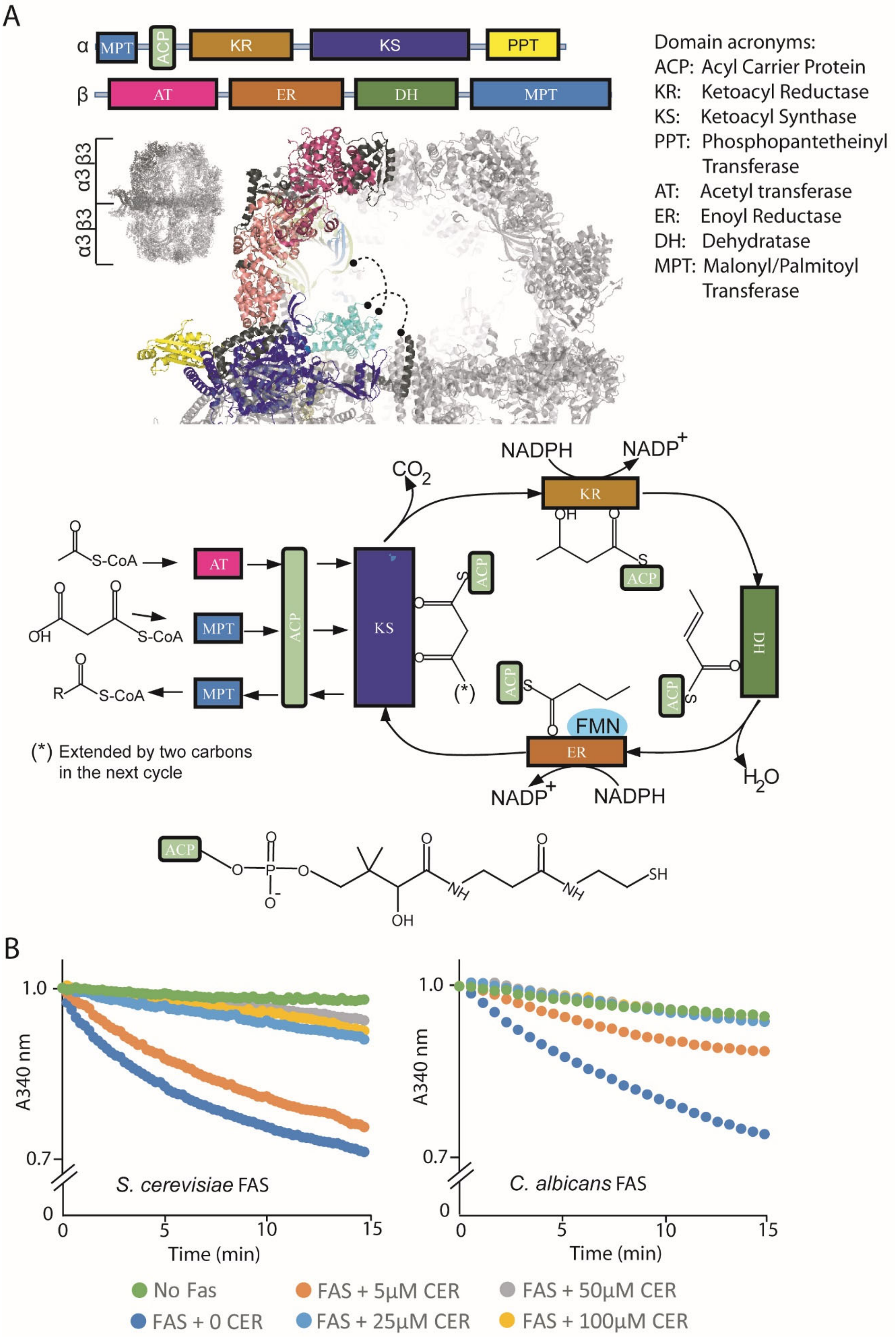

A
α MPT ACP KR KS PPT
β AT ER DH MPT
α3β3
α3β3
Domain acronyms:
ACP: Acyl Carrier Protein
KR: Ketoacyl Reductase
KS: Ketoacyl Synthase
PPT: Phosphopantetheinyl Transferase
AT: Acetyl transferase
ER: Enoyl Reductase
DH: Dehydratase
MPT: Malonyl/Palmitoyl Transferase
NADPH NADP+
CO2
S-CoA
S-CoA
OH
R S-CoA
(*)
(*) Extended by two carbons in the next cycle
FMN
H2O
NADP+ NADPH
SH
B
1.0
0.7
0
A340 nm
Time (min)
S. cerevisiae FAS
C. albicans FAS
No Fas
FAS + 0 CER
FAS + 5μM CER
FAS + 25μM CER
FAS + 50μM CER
FAS + 100μM CER

**Supplementary Figure 1.** (A) Top panel: schematics of the domain organization of *S. cerevisiae* and *C. albicans* FAS. Middle panel: atomic model of the reaction chamber with ACP bound proximal to the KS-domain (PDB: 2PFF[17]). Same color coding as the domain schematics. Bottom panel: schematics of the reaction cycle for palmitoyl-CoA biosynthesis is shown (adopted with modifications from[18]) and chemical structure of the phosphopantetheine arm is shown at the bottom (drawn with ChemSketch, Advanced Chemistry Development, Inc.) (B) Sensitivity of purified *S. cerevisiae* (left) and *C. albicans* (right) FAS to antibiotic cerulenin. The reactions were done in triplicates and a representative activity curve is shown for each condition tested.

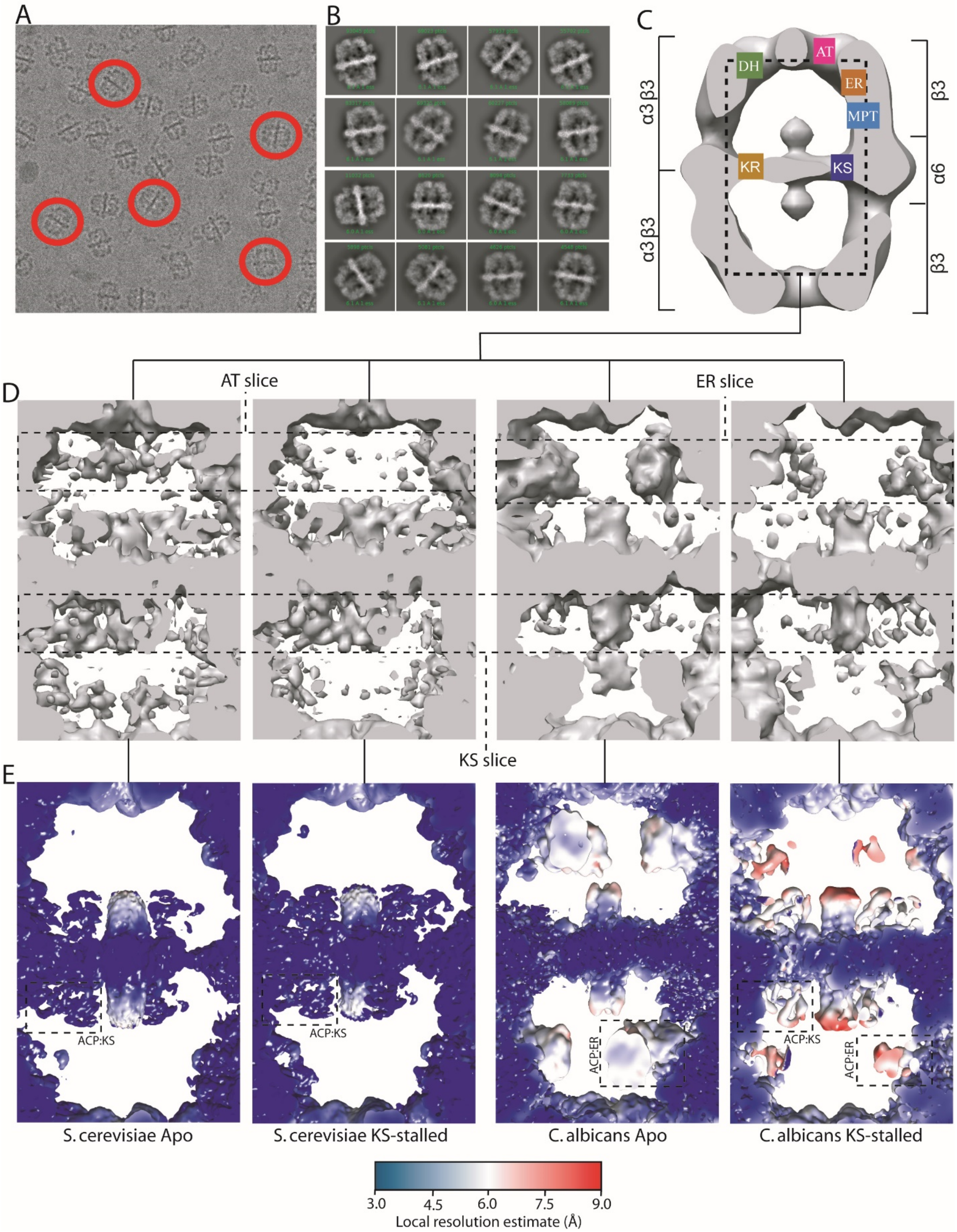

**Supplementary Figure 2.** (A) Example of an aligned and averaged micrograph stack of *S. cerevisiae* FAS particles in vitreous ice. (B) From top to bottom row, 2D classes of *S. cerevisiae* Apo, KS-stalled, and C. albicans Apo and KS-stalled states. (C-E) Scheme for generating *ab initio* ACP densities inside the reaction chamber. Maps are sliced parallel to the long axis (dashed box) to show the interior of the reaction chambers. D3 symmetry applied. (C) initial ACP-less model, (D) *ab initio* reconstructions with experimental particle images. Maps are scaled for each fungal species and shown at identical threshold. Slices through the reaction chamber for ACP at AT, ER, and KS catalytic sites are highlighted with dashed box. (E) high resolution refined maps colored based on local resolution estimate. Representative ACP domains are highlighted with dashed boxes with the name of the interacting catalytic site shown as ACP:catalytic center (*e.g.* ACP:KS).

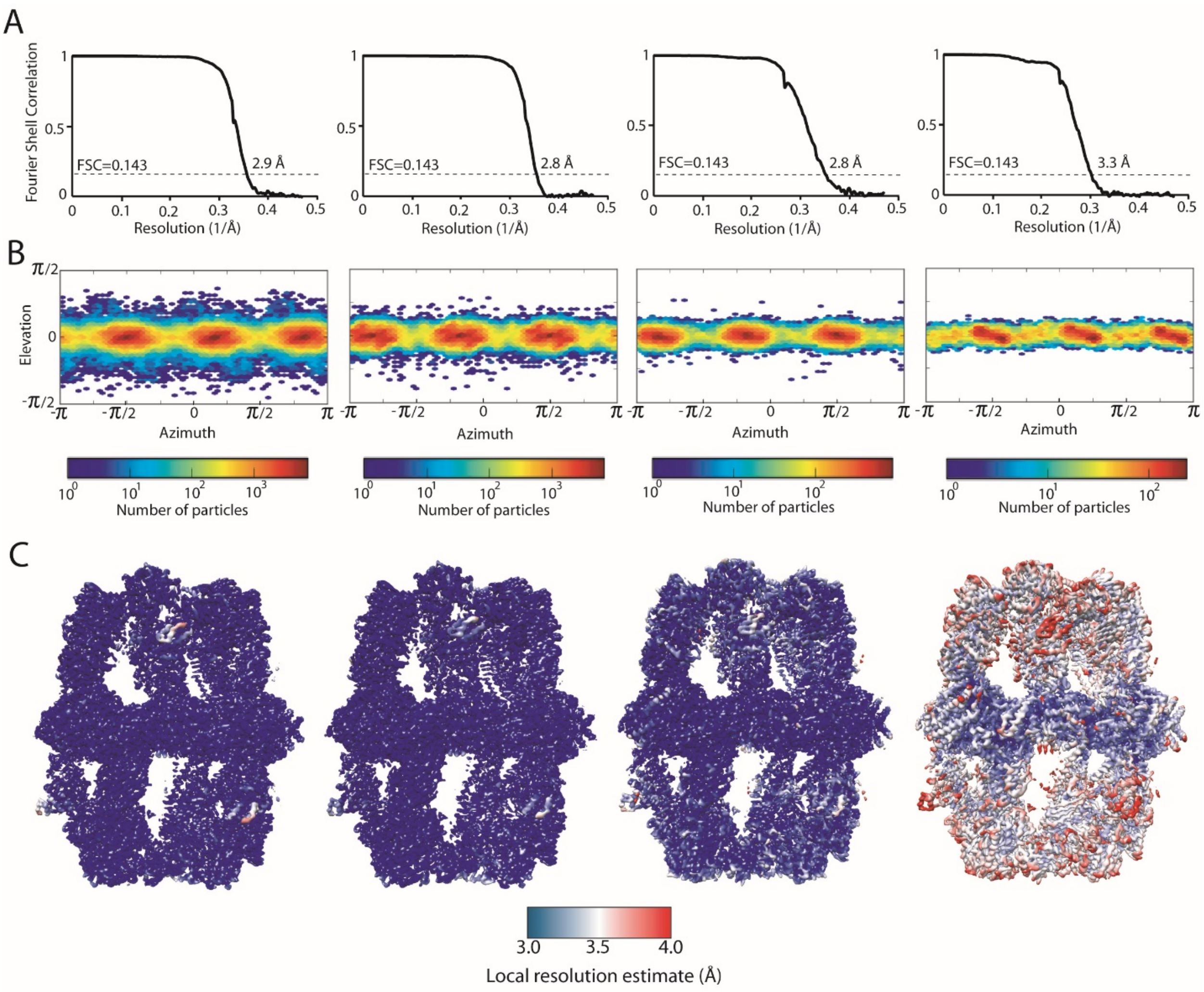

**Supplementary Figure 3.** Quality of the high-resolution maps used in model building. In all panels, left to right: *S. cerevisiae* Apo and KS-stalled, and *C. albicans* Apo and KS-stalled maps. (A) mask-corrected FSC curves, (B) orientation distributions of particle images, and (C) local resolution estimates of the final high-resolution refined maps used in model building. These maps are the same as shown in slices in supplementary Figure 2E but with coloring for different resolution range as indicated.

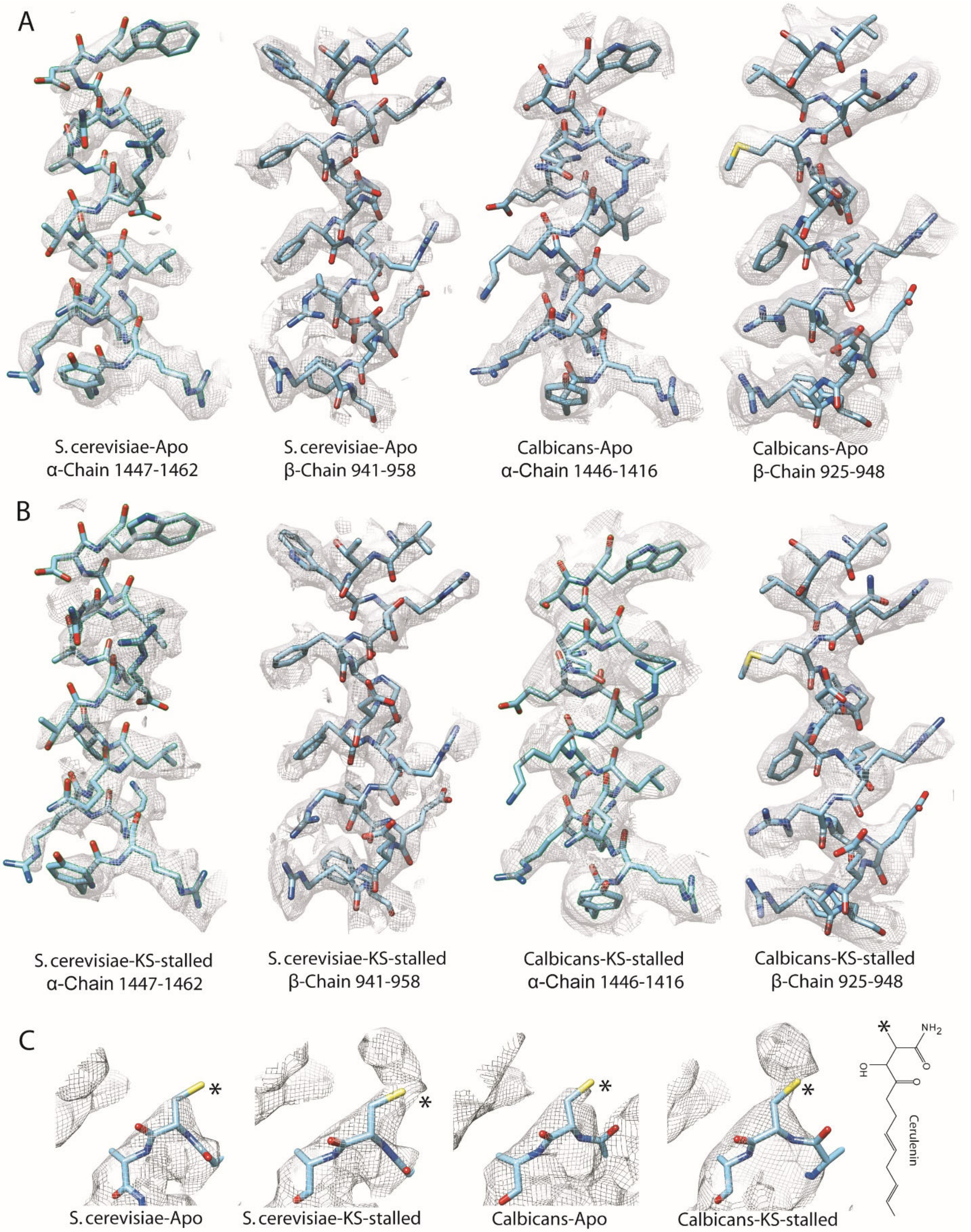

**Supplementary Figure 4.** Quality of model to map fit. Examples of atomic model fit to high resolution refined maps are shown for both α- and β-chains as indicated with respective residue range for (A) Apo and (B) KS-stalled state of each fungal species. (C) Appearance of extra density on -SH functional group (*) of catalytic cysteine of the KS-domain upon inhibition with cerulenin (chemical structure shown to the far right, drawn with ChemSketch, Advanced Chemistry Development, Inc.) for each fungal species. Densities within 5 Å for the catalytic cysteine and the two flanking residues are shown.

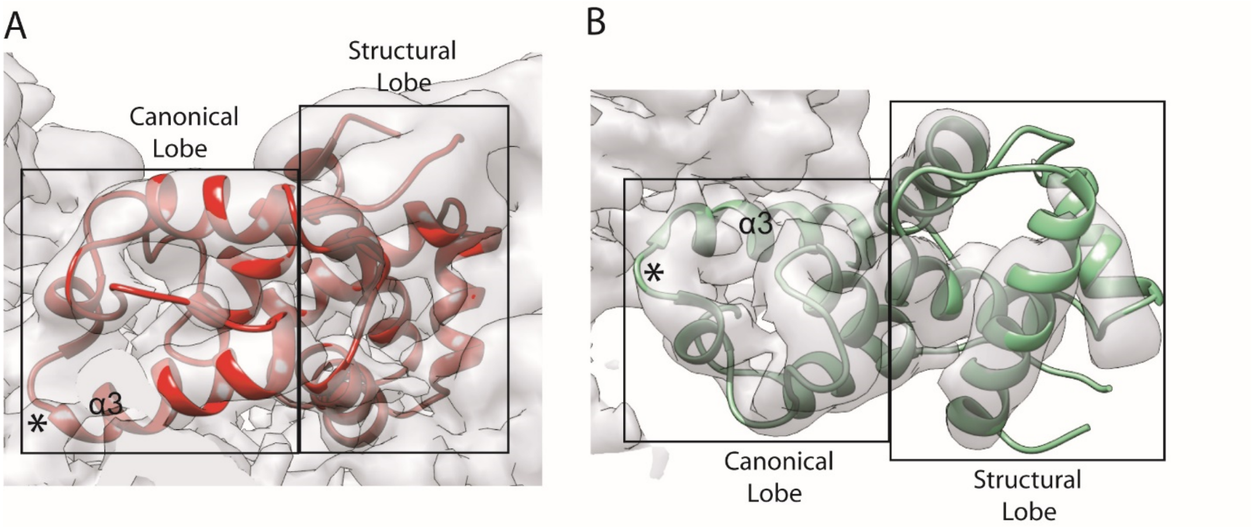

**Supplementary Figure 5.** Fitting of the ACP domains to their corresponding densities near (A) KS- and (B) ER- domains in *S. cerevisiae* and *C. albicans* FAS, respectively. * represents the position of the phosphopantetheine arm.

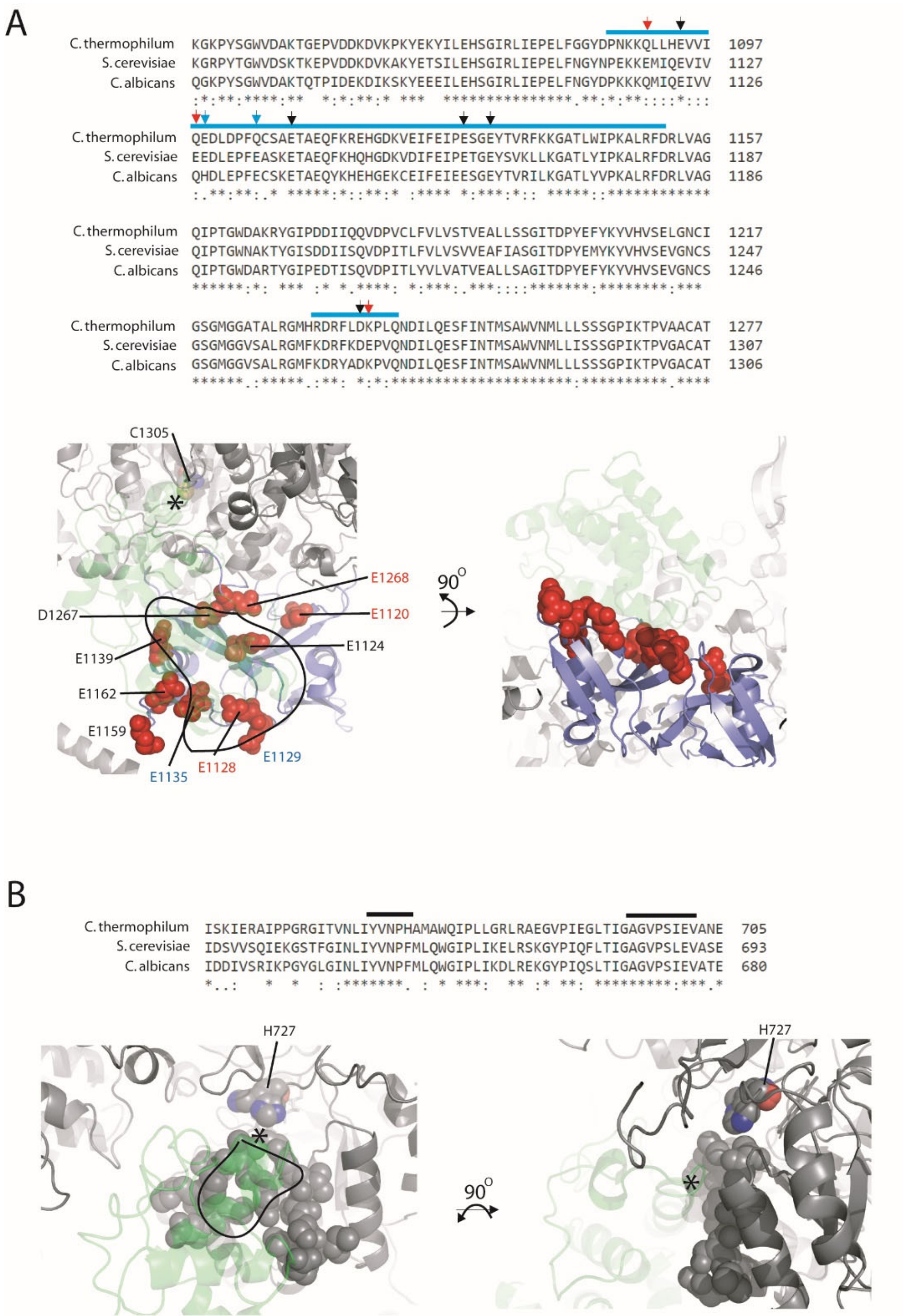

**Supplementary Figure 6.** (A) Negative charged residues in the ACP binding site of the KS-domain for the structural lobe of ACP. Sequence alignment is shown for the three fungal species. The alignment is done on full-length α-chains of each species using Clustal Omega[19] and only the indicated portion is shown. The KS region involved in interaction with ACP structural lobe is highlighted with cyan on top of the sequences and in the structure (bottom panel, model *S. cerevisiae* Apo). ACP is shown as transparent green. Black, blue, and red arrowheads represent KS acidic residues (facing ACP structural lobe) that are present in the three fungal species, only in two (including *S. cerevisiae*), and only in *S. cerevisiae*, respectively. These residues are highlighted on the structure with the same colored residue names (B) ER residues lining the ACP binding sites for the canonical lobe of ACP are highlighted with black bars in the sequence alignment for the three fungal species and shown with sphere in the model of *C. albicans* in the Apo state. Sequence alignment done as above but for the β-chain. ACP is shown as transparent green. * represents the position of the phosphopantetheine arm.

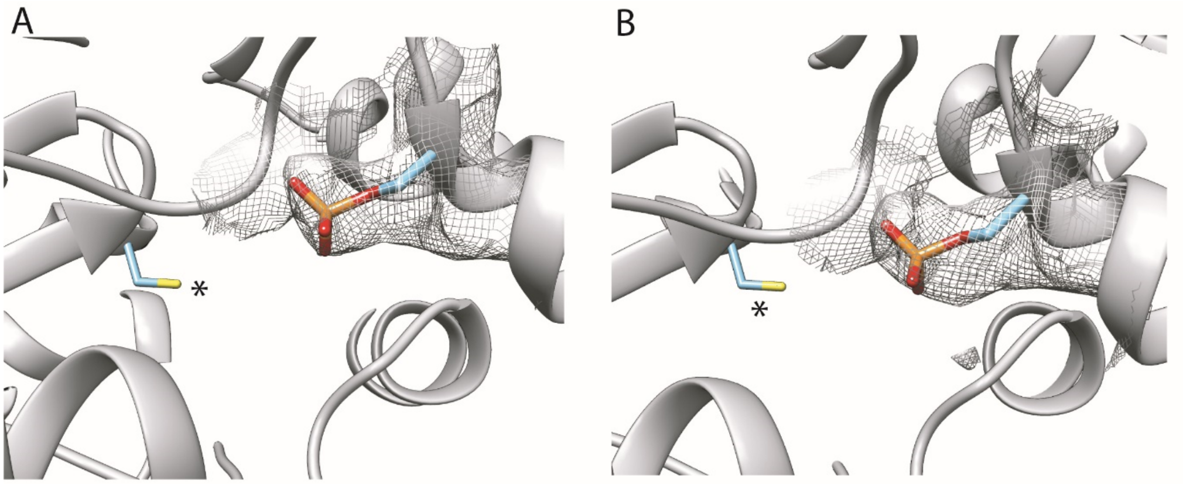

**Supplementary Figure 7.** Density of the phosphopantetheine arm, in the vicinity of the KS-catalytic cavity. cryoEM density maps in (A) Apo and (B) KS-stalled states of *S. cerevisiae* FAS are shown within 5 Å of the partial model of the phosphopantetheine arm at different thresholds to highlight model-to-map fit. * represents the catalytic cysteine of the KS domain.

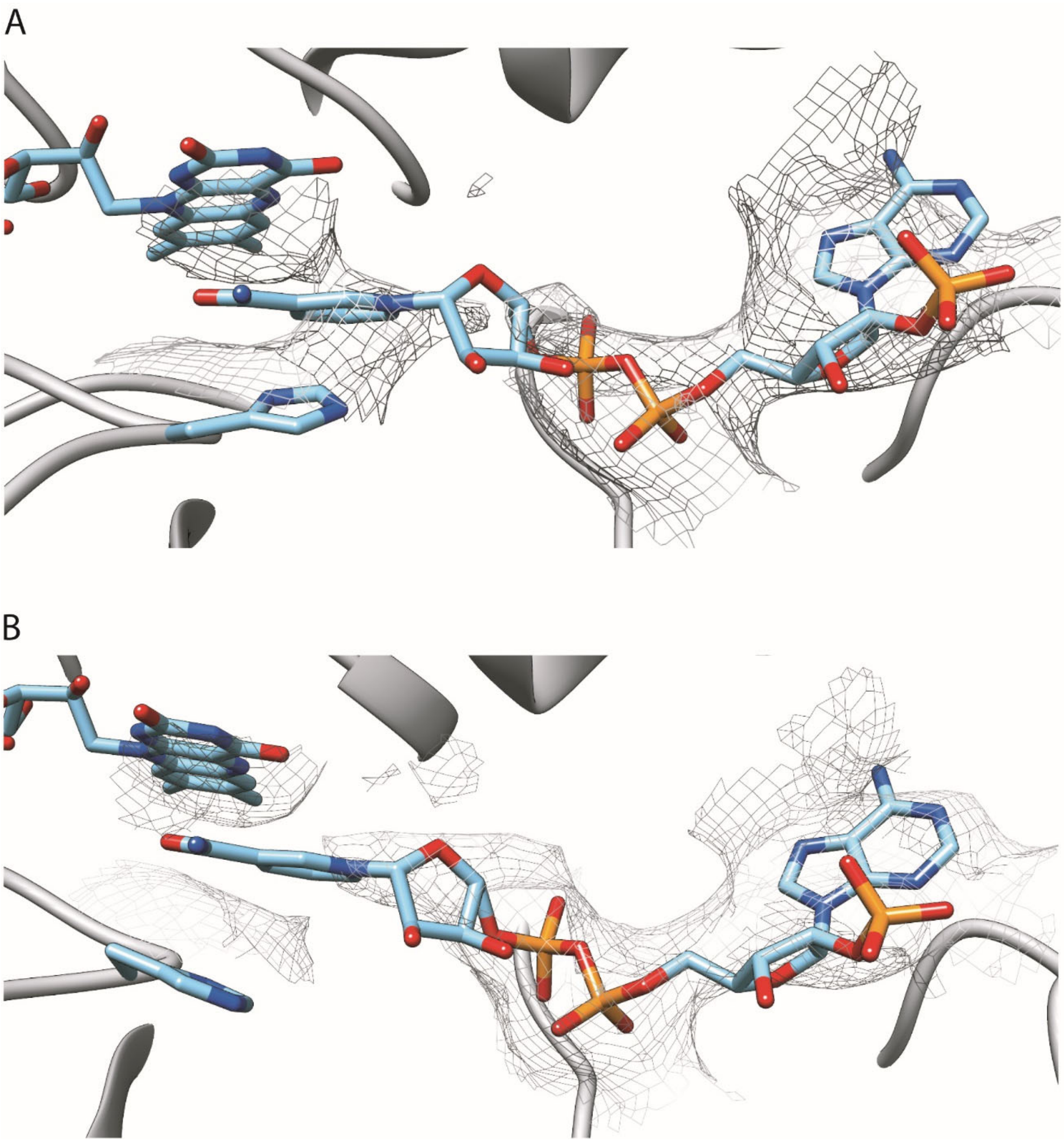

**Supplementary Figure 8.** cryoEM densities of the NADPH molecule in the ER catalytic sites in the KS-stalled states of (A) *C. albicans* and (B) *S. cerevisiae*.
